# Inverse folding of protein complexes with a structure-informed language model enables unsupervised antibody evolution

**DOI:** 10.1101/2023.12.19.572475

**Authors:** Varun R. Shanker, Theodora U.J. Bruun, Brian L. Hie, Peter S. Kim

## Abstract

Large language models trained on sequence information alone are capable of learning high level principles of protein design. However, beyond sequence, the three-dimensional structures of proteins determine their specific function, activity, and evolvability. Here we show that a general protein language model augmented with protein structure backbone coordinates and trained on the inverse folding problem can guide evolution for diverse proteins without needing to explicitly model individual functional tasks. We demonstrate inverse folding to be an effective unsupervised, structure-based sequence optimization strategy that also generalizes to multimeric complexes by implicitly learning features of binding and amino acid epistasis. Using this approach, we screened ∼30 variants of two therapeutic clinical antibodies used to treat SARS-CoV-2 infection and achieved up to 26-fold improvement in neutralization and 37-fold improvement in affinity against antibody-escaped viral variants-of-concern BQ.1.1 and XBB.1.5, respectively. In addition to substantial overall improvements in protein function, we find inverse folding performs with leading experimental success rates among other reported machine learning-guided directed evolution methods, without requiring any task-specific training data.

## Introduction

Evolution generates diverse proteins at the level of biological sequences by exploring a vast search space of potential mutations and acquiring those that improve fitness. However, it is the three-dimensional structure encoded by these sequences that ultimately determines the function and activity of a protein. Consequently, as proteins accumulate mutations, they undergo corresponding structural changes, which in turn facilitate functional adaptations^1^.

In the laboratory, this tendency for greater sequence change to cause structural divergence poses a major challenge to engineering better proteins via a stepwise evolutionary process. Mutations added in sequential rounds of artificial evolution are increasingly likely to destabilize the structure and therefore diminish the protein’s evolvability^2^. Identifying beneficial mutations is further challenged by the fact that almost all mutations to a prototypical protein are deleterious, or at best neutral, and only a rare subset are beneficial on its fitness landscape^3–8^. In total, these phenomena can often reduce the evolutionarily accessible paths and make evolution more susceptible to local fitness optima^9,10^, further complicating attempts to increase fitness.

To address both the structural constraints of protein design and the high dimensionality of the mutational search space, we utilized a general protein language model augmented with structural information and trained across millions of non-redundant single sequence-structure pairs on the inverse folding objective^11^. Most simply, the inverse folding problem considers the task opposite of that performed by many of the recent powerful structure-prediction tools, including AlphaFold and ESMFold^12,13^: recovery of a protein’s native sequence, given its three-dimensional backbone coordinates (**Figure 1a**). This is accomplished by predicting the identity of an amino acid given both the preceding amino acid sequence (referred to as autoregressive modeling) and the entire structure’s backbone coordinates (**Methods**). Thus, sequences assigned high likelihood scores by the inverse folding language model are expected to fold into the backbone of the input structure with high confidence (**Figure 1b**).

**Figure 1:**
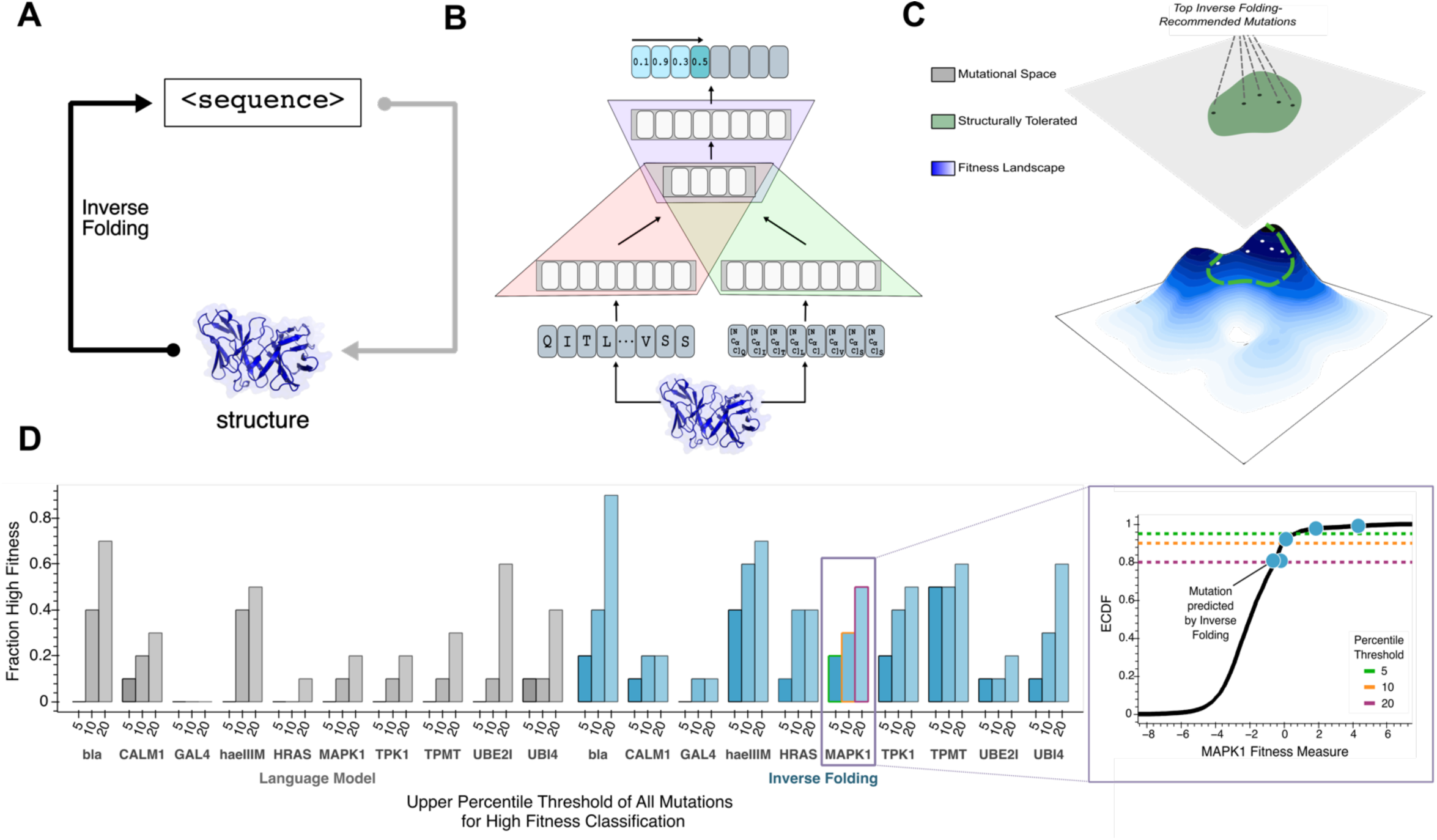
Guiding evolution of diverse proteins via inverse folding. **(A)** The inverse folding problem refers to the prediction of a protein’s native amino acid sequence, given its three-dimensional backbone structure, which is conceptually analogous to the opposite problem solved by structure prediction tools like AlphaFold^12^. **(B)** A hybrid autoregressive model^11^ integrates amino acid values and backbone structural information to evaluate the joint likelihood over all positions in a sequence. Amino acids from the protein sequence are tokenized (red), combined with geometric features extracted from a structural encoder (green), and modeled with an encoder-decoder transformer (purple). Sequences assigned high likelihoods by the model represent high confidence in folding into the input backbone structure. **(C)** Our structure-guided framework for protein design indirectly explores the underlying fitness landscape, without modeling a specific definition of fitness or requiring any task-specific training data, by constraining the search space to regions where the backbone fold preserved. **(D)** High fitness sensitivity analysis reveals that multimodal input improves language model performance compared to sequence-only input across 10 proteins from diverse protein families (left). ‘Fraction High fitness’ is the fraction of the top ten single amino acid substitutions recommended by each model that are ranked in the top indicated percentile of all experimentally screened variants. A representative plot (right) demonstrates this metric for assessing enrichment of high-fitness MAPK1 mutations, with successfully predicted mutations highlighted (blue) on the empirical cumulative density function (ECDF) of the experimental data (black). The three different thresholds, as defined by percentiles, are also shown as dashed lines. Inverse folding predictions are more enriched, on average, for high fitness variants across various tested thresholds for high fitness classification. bla, Beta-lactamase TEM; CALM1, Calmodulin-1; haeIIIM, Type II methyltransferase M.HaeIII; HRAS, GTPase HRas; MAPK1, Mitogen-activated protein kinase; TMPT, Thiopurine S-methyltransferase; TPK1, Thiamin pyrophosphokinase 1; UBI4, Polyubiquitin; UBE2I, SUMO-conjugating enzyme UBC9

Our inverse folding framework for protein design does not model an explicit protein function or definition of protein fitness. Rather, using a structure-guided paradigm, we indirectly explore the underlying fitness landscape by focusing exploration to regions where the backbone fold of the protein is preserved. We hypothesize constraining evolution to regimes of high inverse folding likelihood can serve as an effective prior for high-fitness variants, and thereby improve the efficiency of evolution (**Figure 1c**).

We reasoned that this approach may be particularly valuable for the evolution of human antibodies, which are used clinically to treat a broad range of diseases^14^. Antibodies are used therapeutically to bind to a target antigen mediating pathogenesis, and modify or disrupt its function^15^. A central concept of this study is to use the complete structure of the antibody-antigen complex to guide evolution. By conditioning the inverse folding model on the entire antibody-antigen complex, we sought to enable the discovery of mutations that preserve or enhance the stability of the entire complex, and thus that improve antibody function.

Indeed, we show that as an unsupervised machine learning-guided evolution strategy, inverse folding is capable of identifying high fitness mutations across several protein families and tasks, performing better than sequence-only methods. We found that inverse folding generalizes to protein complexes with improved antibody variant prediction when antigen structural information is also included as input. To demonstrate the practical utility of this method, we improved the potency of mature, clinical SARS-CoV-2 monoclonal antibody therapies, in a low-throughput setting, against both their original viral target as well as viral escape variants that reduced their efficacy, namely variants-of-concern (VOC) BQ.1.1 and XBB.1.5. We achieved up to 26-fold improvement in the neutralization potency of Ly-1404 (Bebtelovimab) against BQ.1.1, and 11-fold for SA58, testing only a total of 31 and 25 antibody variants, respectively. We also achieved 27-fold improvement in affinity against BQ.1.1 and 37-fold improvement in affinity against XBB.1.5. Notably, all experimentally tested combinations of inverse folding-recommended mutations showed improved activity, with many designs comprising multiple synergistic mutations. With our approach, we report experimental success rates that surpass all previous machine learning-guided protein evolution methods^8,16–28^, including those based on supervision with task-specific training data. These findings highlight the advantage of an unsupervised, structure-based paradigm to identify efficient evolutionary trajectories.

## Results

### Inverse folding enriches sequence exploration for high function protein variants across diverse tasks

We evaluated whether inverse folding can be used to guide protein evolution, without needing to explicitly model specific functional tasks, by assessing its ability to identify mutations resulting in high levels of protein activity for a desired functional property, or fitness measure. Accordingly, for 10 proteins from diverse families among four organisms, and with functions ranging from enzyme catalysis (TPMT) to oncogenesis (HRAS) to transcriptional regulation (GAL4), we used inverse folding likelihoods to score variants profiled in large datasets from deep mutational scanning experiments^29–38^ against a target backbone of the wild-type protein^39–48^ (**Methods**, **Supplementary Table 1**).

From the thousands of tested variants for each of the 10 proteins, we identified numerous with experimentally determined protein activities ranking in the top percentiles of the entire screen within just the set of top ten inverse-folding predictions (**Figure 1d**). Our analysis also demonstrates that conditioning on structural information serves to improve predictive capabilities of protein language models as we successfully identified mutations in the top fifth percentile for 9 out of the 10 proteins using inverse folding compared to just 2 proteins using a state-of-the-art general protein language model trained only on sequence information and specifically for variant prediction (ESM-1v)^49^ (**Figure 1d**). This improvement in prediction also holds with increasingly relaxed thresholds for classification as high-fitness variants.

These results suggest that inverse folding offers a promising alternative to brute force experimental searches for beneficial mutations. Notably, some of the top mutations predicted by inverse folding are also the same ones recovered from exhaustive experimental exploration. For example, for restriction enzyme haeIIIM, variant Q18E is recommended within the top five inverse folding predictions and experimentally ranks as the second-best substitution (and > 5 standard deviations above the mean) out of the nearly 2000 substitutions screened to the endonuclease^38^. Another key advantage of our task-independent framework, in addition to being broadly applicable across diverse proteins, is the ability to improve a single protein for multiple desired properties without needing to develop specialized high-throughput assays to screen each independently. From just the top 10 inverse folding predictions for MAPK1, we identify substitutions Q105M and Y64D, which are experimentally shown to confer resistance to two different oncogenic-targeting MAPK1 kinase inhibitors^32^.

### Inverse folding is a state-of-the-art zero-shot mutational effect predictor for antibodies

To analyze the effectiveness of augmenting a general protein language model with structural information, specifically for antibody variant prediction, we compared the inverse folding likelihoods of sequences across entire mutational landscapes against the corresponding experimental fitness values from three existing mutagenesis datasets. The first two of the datasets profile the scFv equilibrium dissociation constants (*K*_D_) of all possible evolutionary intermediates between the inferred germline and somatic sequence of naturally affinity-matured influenza broadly neutralizing antibodies (bnAbs) CR9114 and CR6261, which bind the conserved stem epitope of influenza surface protein hemagglutinin (HA)^50^. For both bnAbs, only mutations in the heavy chain, which is responsible for antigen binding, were characterized. The profiled mutational landscape of CR9114 includes all possible combinations of 16 substitutions while that of CR6261 includes all possible combinations of 11 substitutions, totaling 2^16^ = 65,536 and 2^11^ = 2,048 variant antibody sequences respectively. Each of these libraries were screened for binding against two distinct influenza HA subtypes (H1 and H3 for CR9114 and H1 and H9 for CR6261). The third dataset assesses the effects of all possible single amino acid substitutions with a deep mutational scan profiling 4,275 mutations in the variable regions for both heavy chain (VH) and light chain (VL) of antibody G6.31 to binding with its ligand, vascular endothelial growth factor A (VEGF-A)^51^.

For each dataset, we computed the Spearman correlation between the log likelihood estimated by the inverse folding model and the experimentally determined binding measure for a given antigen, across all sequences in the mutational library. We scored the inverse folding likelihood of each candidate sequence in the library using the backbone coordinates of a structure with the mature antibody bound to its target antigen^52–54^.

Across all five experimental binding datasets, we found that inverse folding performs better than both a sequence-only language model, ESM-1v^49^, and a site-independent model of mutational frequency curated with extensive antibody sequence alignments, abYsis^55^. In nearly all experimental scenarios, supplementing sequence information with the backbone coordinates of the antibody alone, without providing antigen information, as input to inverse folding is sufficient to outperform other sequence-only methods. A notable feature of the autoregressive architecture is that it computes the joint likelihood over all positions in a sequence, making it well-suited to score combinatorial sequence changes. We find that inverse folding can capture complex epistatic interactions, or potential interdependence among individual amino acids, as it performs well on the CR9114 and CR6261 libraries composed of sequences with multiple mutations (**Figure 2a,b**).

**Figure 2:**
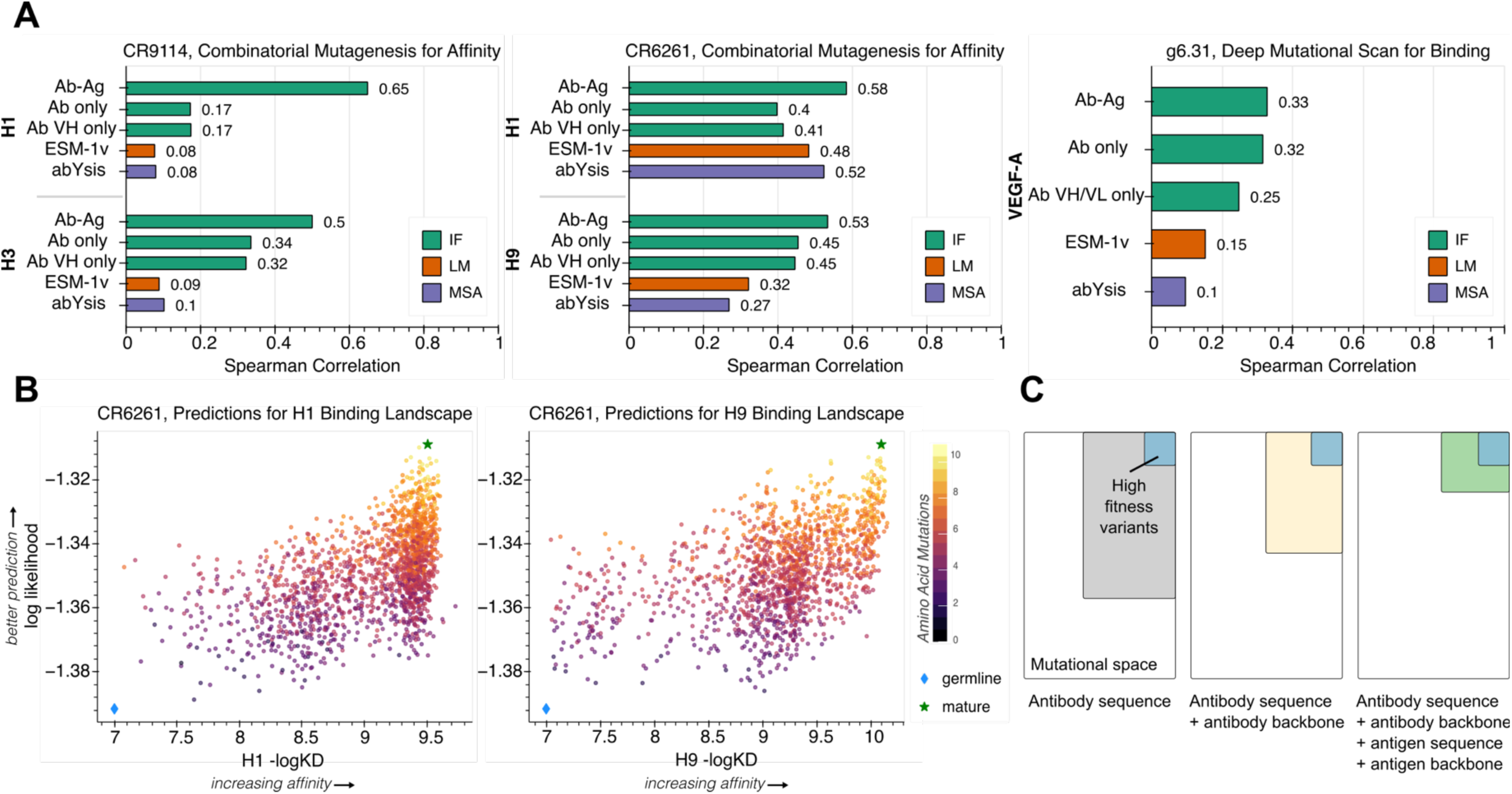
Inverse folding of antibody-antigen complexes resolves mutational landscapes by implicitly learning features of binding and protein epistasis. **(A)** Spearman correlation using inverse folding as well as sequence-based modeling approaches ESM-v^49^ and abYsis^55^ reported for three antibodies screened with corresponding influenza A HA subtypes H1, H3, and H9. Bars are colored by the type of model used: IF, Inverse Folding (green); LM, Language Model (orange); and MSA, Multiple Sequence Alignment (purple). Inverse folding was evaluated in three different settings: i) providing the entire antibody variable region and antigen complex (Ab-Ag) ii) providing only the antibody variable region (Ab only), and iii) providing only the single antibody variable region of the chain responsible for binding or being mutated (Ab VH only or Ab VH/VL only). Inverse folding implicitly learns features of binding and protein epistasis. For example, when scoring combinatorial mutations to CR9114 against H1, we find that the model has much higher performance (Spearman π = 0.65 for H1, 0.5 for H3) than a masked language model ESM-1v (Spearman π = 0.08 for H1, 0.09 for H3) and a site-independent, alignment-based model abYsis (Spearman π = 0.08 for H1, 0.1 for H3). This performance improvement is also consistent across the other combinatorial landscapes tested. **(B)** Scatter plots showing inverse folding predictions against experimentally determined dissociation constants of CR6261 against HA-H1(left) and HA-H9 (right). The germline and mature sequences are highlighted on all plots as indicated in the legend. For visualization, all scatter plots omit points on the lower limit of quantitation. Further analysis of assay limit on predictive performance is shown in **Supplementary** Figure 2. **(C)** Conceptual schematic representation of protein language performance improvements with improved priors. Providing sequence and structural information of both the antibody and antigen enables inverse folding to most efficiently identify complex destabilizing mutations and enrich for high fitness antibody variants.

We achieved the greatest improvement in performance on all five experimental screens by incorporating the structure of both the antibody and antigen (**Figure 2a**), indicating that the inverse folding model can implicitly learn features of binding (**Figure 2c**). This result is particularly significant, given that the inverse folding model is only trained on single-chain protein structures, while the antibody-antigen complexes we use as inputs are composed of either three (G6.31) or four (CR9114, CR6261) protein chains. The most substantial contribution of antigen information is observed in the case of CR9114-H1, for which the correlation increases from 0.17 with only antibody information to 0.65 with sequence and backbone coordinates of the entire complex.

Remarkably, we could still predict effects of mutations on binding for a cross-reactive antibody while using a different antigen as input to the model. (**Figure 2a,b**). Despite using a complex with HA from H5N1 influenza as input to score CR9114 variants, we obtain correlations of 0.65 and 0.50 with experimental binding data for H1 and H3, respectively. This is particularly striking since, for example, H5 and H1 only share 63% sequence identify across both HA subunits (**Supplementary** Figure 3). This same cross-reactive predictive capability is observed for CR6261, which is tested experimentally against H1 and H9 while we use an input structure with HA from 1918 H1N1 influenza (**Figure 2a**). Although inverse folding cannot learn explicit chemical rules of binding (e.g., hydrogen bonding or disulfide bridge formation) since it does not have access to amino acid side chain atomic coordinates, these results suggest that structural principles like interface packing or potential steric interference are not only implicitly accessible from residue identities, but are also informative for binding prediction.

Our model’s top recommended mutations are made independently of a specific definition of fitness; they simply represent a set of variants with a high likelihood of folding into the input backbone structure. Therefore, our model’s recommendations may also help identify mutations that improve other useful biochemical properties beyond affinity. Impressively, for example, the top inverse folding-recommended mutation to the VL of G6.31 is F83A, which was identified in the original screening study^51^ to be particularly interesting as it confers a three-fold increase in VEGF-A binding affinity and a 5°C improvement in melting temperature, despite being 25Å from the antigen and in the antibody framework region. It was determined that the VL F83A substitution induces more compact packing and the site serves as a conformational switch that affects biological activity at the antibody-antigen interface by modulating both interdomain and elbow angle dynamics^51^.

### Engineering therapeutic antibodies for increased potency and resilience

Finally, we aimed to assess if the structure-augmented language model’s predictive capabilities could not only resolve trends on large sets of experimental data, but also enable efficient and successful directed evolution campaigns while testing only a small number (on the order of tens) of variants. To do so, we considered the task of improving the potency and resilience (effectiveness against a virus as it mutates over time) of two mature, clinical monoclonal antibody therapies.

- Ly-1404 (Bebtelovimab) was isolated from a COVID-19 convalescent donor and binds to the receptor binding domain (RBD) of the SARS-CoV-2 Spike protein^56^. It was approved by the U.S. F.D.A. on February 11, 2022 given its activity against both the original Wuhan and Omicron SARS-CoV-2 variants and was the last remaining approved monoclonal antibody therapy withstanding against viral evolution^57^ until its discontinuation on November 30, 2023 due to antibody evasion by VOC BQ.1.1.^58^
- SA58 (BD55-5840) was isolated from a vaccinated individual and is one of two RBD-targeting neutralizing antibodies (NAb) in a rationally developed antibody cocktail. SA58 alone retained efficacy against all Omicron subvariants, including *in vivo* protection against BA.5^59,60^ and was shown to be effective as a post-exposure prophylaxis in a clinical study^61^.

For both antibody engineering campaigns, we used the inverse folding language model to compute likelihoods of all ∼4,300 possible single-residue substitutions in the VH or VL regions of the antibody. In the first round of evolution, we selected only the top ten predictions at unique residues in each chain for experimental validation. An important practical benefit of our method is the ability to optimize against measures of fitness most relevant to the protein’s downstream function, rather than being limited to indirect and less accurate surrogate measures that are more amenable to high-throughput screening^4,16^. We leverage this advantage to directly evolve these antibodies for their ability to more potently neutralize SARS-CoV-2 pseudotyped lentivirus.

Variants recommended by the inverse folding language model were assessed by comparing the half-maximal inhibitory concentration (IC_50_) relative to the wild-type antibody. Remarkably, although we chose to only test 20 single-site substitutions for each of the two clinical monoclonal antibody therapies, approximately one-third of them improved neutralizing potency. Notably, several of these variants improve neutralization IC_50_ by approximately 2-fold with just a single amino acid change (**Figure 3a**, **Supplementary Data 1**).

**Figure 3.**
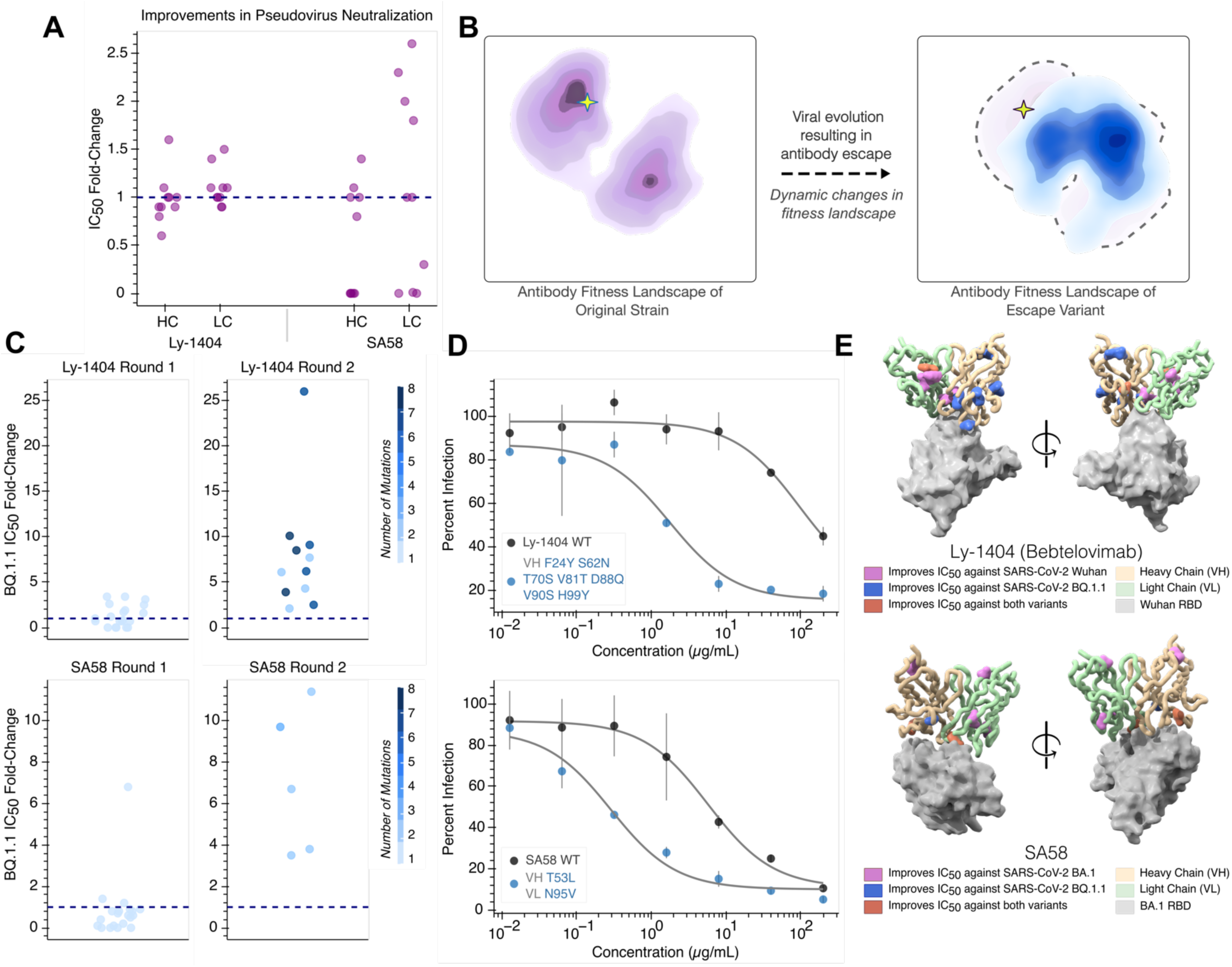
Inverse folding-guided evolution of antibodies improves neutralization potency and resilience. **(A)** Each point represents the fold-change in IC50 of pseudovirus neutralization for antibody variants with single amino acid mutations. Antibodies are tested against the viral strain represented in the input structure (Ly1404-Wuhan, SA58-BA.1 Omicron). A dashed line is shown at fold-change of 1 corresponding to no change. 35% of Ly-1404 variants and 30% of SA58 variants improved antibody potency (defined as 1.1-fold or higher improvement in IC50 compared to wild-type). Among this subset of beneficial mutations, we identify single amino acid mutations that provide a 1.6-fold improvement in Ly-1404 IC50 and a 2.6-fold improvement in SA58 IC50. **(B)** Conceptual representation of viral evolution. Selection for immune evasion drives antibody escape, which fundamentally represents a dynamic change in the underlying fitness landscape for the antibody. This antigenic drift displaces a potent antibody from a peak on the previous fitness landscape (left) to a new starting point at lower activity (right). **(C)** Strip plots visualizing antibody evolution across two rounds. Each point shows the corresponding fold-change in IC_50_ of pseudovirus neutralization for a designed variant and is colored according to the number of mutations it has (1-8). Consistent with preserving backbone fold, all 55 designed variants across both antibody evolutionary campaigns could be expressed. All round 1 variants are only composed of only single amino acid changes while beneficial mutations are combined in round 2. All round 2 variants have improved neutralization activity compared to their respective wild-type antibody (dotted line). **(D)** Pseudovirus neutralization curves are shown for the most potent evolved antibody variant, consisting of mutations annotated to the left. The top Ly-1404 variant, bearing seven amino acid substitutions in VH, achieves a 26-fold improvement in neutralization against BQ.1.1 (top). The top SA58 variant, bearing single amino acid mutations in both VH and VL, achieves an 11-fold improvement in neutralization against BQ.1.1 (bottom). **(E)** Residues at which mutations improve neutralization against either the structure-encoded strain, BQ.1.1, or both viral strains are highlighted with spheres for antibodies Ly-1404 (PDB 7MMO) and SA58 (PDB 7Y0W). Notably, beneficial mutations are identified both within the binding interface as well distal to the antigen. Neutralization enhancing mutations are labeled in **Supplementary** Figure 6.

Prompted by recent evidence showing that conservation of the overall RBD structure is robust to SARS-CoV-2 evolution^62^, we next sought to determine whether we could also evolve the previously mature antibodies against SARS-CoV-2 BQ.1.1, the variant responsible for diminished therapeutic efficacy. Although the antibodies were previously effective, a change in antigen conceptually represents a fundamental shift in the underlying fitness landscape (**Figure 3b**). From the same set of 20 single amino acid substitutions to Ly-1404, we found that nearly half improve neutralization of variant BQ.1.1. In addition to a high success rate, we also found multiple of these mutations provided a large magnitude of improvement. Several single amino acid substitutions to Ly-1404 individually result in over a 3-fold improvement while the most beneficial mutation to SA58 results in a nearly 7-fold improvement (**Figure 3c**).

Taken together, approximately two-third and one-third of tested single amino acid substitutions to Ly-1404 and SA58, respectively, were beneficial for neutralization of either the original strain or BQ.1.1. These results reinforce that, despite all being predicted to have the same backbone fold, inverse folding variants feature functional diversity and can be used for distinct notions of protein fitness. Interestingly, for both antibodies, the most beneficial mutation, is not shared by the each of the strains tested (**Supplementary** Figure 4).

A common challenge in directed evolution is contending with the combinatorial explosion of possible sequences which emerges from trying to combine a set of individually beneficial mutations. In the second round of evolution, we simply use the inverse folding model again to acquire up to five top-scoring unique combinations of mutations to each antibody chain (**Methods**). Notably, across both evolutionary trajectories, all 15 antibody designs with multiple mutations have IC_50_ values better than wild-type, with many designs showing synergistic effects upon combination. For example, just a single amino acid mutation in each of the two chains of SA58 leads to over an 11-fold improvement (**Figure 3c,d**). Similarly, the most potent evolved design of Ly-1404 is a combination of seven of the eight beneficial single amino acid substitution to the VH and improves neutralization 26-fold (**Figure 3d**). Critically, these improvements to neutralizing potency against BQ.1.1 do not sacrifice potency against the original strains. We found that the top SA58 design against BQ.1.1 after the second round of evolution also improves BA.1 neutralization nearly 3-fold (**Supplementary Data 1**).

### Additional characterization of evolved antibodies

To further characterize the basis for enhanced neutralization of SARS-CoV-2 VOC BQ.1.1, we tested the binding affinity of all variant antibodies to RBD as bivalent IgG using biolayer interferometry (BLI) to obtain the apparent dissociation constant (*K*_D,app_). For Ly-1404, all 23 variants with improved neutralization also have improved binding affinity up to ∼27-fold. Interestingly, we found four additional inverse folding-recommended mutations, which were neutral or deleterious to neutralization, also improved binding affinity. Across all variants there is a Spearman correlation of 0.47 between fold-change in IC_50_ and fold-change in *K*_D,app_ (**Figure 4a**).

**Figure 4:**
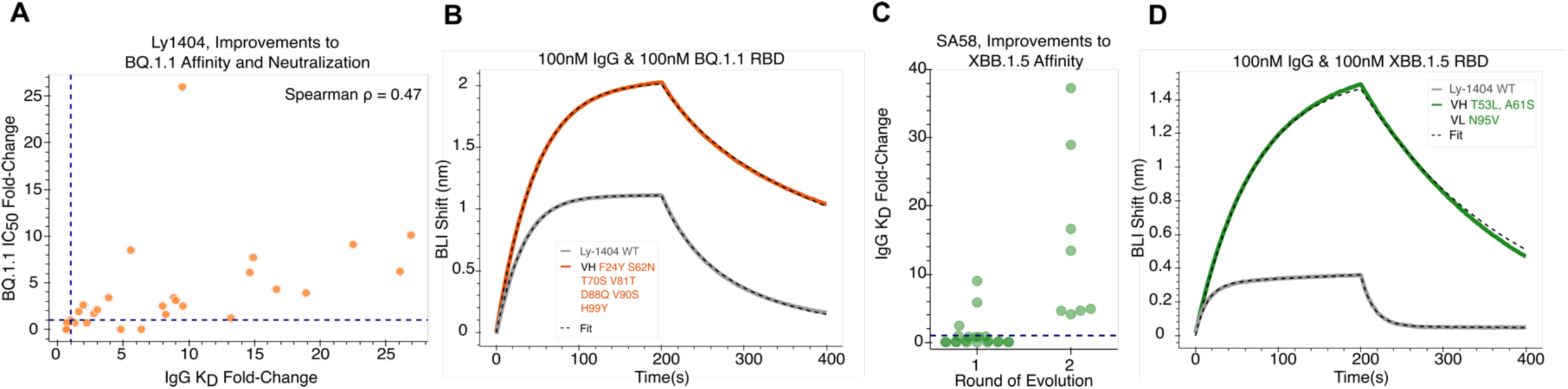
Antibodies evolved for high potency also exhibit improved affinity. **(A)** Ly-1404 antibody variants show a Spearman correlation of 0.47 between apparent affinity fold-change and potency fold-change. Improved affinity is observed to be necessary but not sufficient for improved neutralization activity. Four variants exhibit improved affinity but do not enhance neutralization. All variants with improved neutralization also display improved affinity. The top inverse folding Ly-1404 design with a 27-fold improvement in neutralization has a 9.5-fold improvement in affinity to BQ.1.1 RBD, as measured using BLI. **(C)** SA58 antibodies evolved for improved potency against BQ.1.1 also exhibit improved affinity against VOC XBB.1.5, up to 37-fold. **(B, D)** Representative traces of BLI binding assays for Ly-1404 and SA58 variants with improved affinity.

We similarly screened the SA58 variants for binding to the RBD of BQ.1.1. However, since the *K*_D_ of the wildtype antibody as IgG was already sub-picomolar, further improvements to binding were below the limit of quantitation and indistinguishable using this measure. Given this strong binding affinity of wildtype SA58 to BQ.1.1 RBD, we also screened this same set of variants against emerging VOC XBB.1.5 and observe improvements in *K*_D,app_ up to 37-fold (**Figure 4c**).

By testing several top affinity-matured designs in a polyspecificity assay, we also confirmed that improvements in binding are not mediated by generalized enhancements of non-specific interactions (**Supplementary** Figure 5a). In this assay, we observed no substantial changes in off-target binding of the evolved antibodies to membrane soluble proteins, particularly within a therapeutically viable range (as defined by controls of clinically approved antibodies with recorded high and low polyspecificity). Furthermore, we found no correlation between fold-change in polyspecificity and affinity fold-change (**Supplementary** Figure 5b).

### Analysis of evolutionary exploration

Confronted by the large number of possible mutations, traditional experimental-based methods for antibody affinity maturation often restrict the mutational search space to only a few regions of the antibody. Specifically, binding optimization efforts are typically focused within the complementarity determining regions (CDR), which are hotspots for natural somatic hypermutation. However, using our unbiased approach to consider all regions of the variable domain allows for many discoveries that may be less intuitive to a rational designer. For example, the most beneficial substitutions to Ly-1404, VH F24Y and VH V90S, are located within framework regions and positioned distally from the binding interface (**Supplementary** Figure 6**, Supplementary Table 2**). Interestingly, they both improve neutralization of BQ.1.1 by over 3-fold and are not deleterious to Wuhan neutralization. In other cases, inverse folding also successfully predicts beneficial substitutions using residues rarely observed among human antibody sequences. Substitution VL N95V in SA58, which improves neutralization approximately 7-fold against BQ.1.1, is mediated by the incorporation of a valine observed in only 0.7% of human antibody sequences at that position and enhances antibody-antigen contact. While inverse folding is capable of successfully making novel predictions, in some instances it also does suggest reverting residues to ones frequently selected for in natural somatic hypermutation. Mutation VL F51Y in Ly-1404 changes a phenylalanine observed in just 5% of sequences to a tyrosine observed in 86% of sequences. However, this variant results in no change to Wuhan neutralization. Overall, these results highlight the novelty and value in augmenting a language model with structural information to evolve antibodies and proteins complexes.

## Discussion

The discovery of mutations that improve protein function is inherently challenging due to the large sequence search space and complex rules that govern the relationship between sequence and function, such as stability or environmental selection pressures. We show that a general inverse folding protein language model informed with the sequence and backbone structural coordinates of a protein can considerably improve directed evolution efforts by serving as an improved prior compared to sequence-only deep learning methods. Importantly, we highlight that inverse folding can interrogate protein fitness landscapes indirectly, without needing to explicitly model individual functional tasks or properties, making it broadly applicable to proteins across diverse settings ranging from enzyme catalysis to antibiotic and chemotherapy resistance (**Figure 1d**). We also demonstrate inverse folding generalizes to multimeric proteins, despite being trained only on single-chain proteins, through its ability to implicitly learn features of binding. This result is particularly remarkable considering inverse folding has no access to amino acid side chain atoms, coordinates, or bond information.

Equipped with these capabilities, we use inverse folding to evolve clinical therapeutic antibodies and identify several mutations which act synergistically to improve antibody potency and resilience against emerging variants of concern. In the context of pandemics and emergency-use situations, where monoclonal antibody therapies are limited in supply and vulnerable to resistance from viral evolution, the ability to rapidly make improvements in potency with a general method could have major clinical and economic implications.

In comparison to fourteen other promising machine learning-guided protein design methods^8,16–28^, we find that inverse folding has the strongest performance to date, even without requiring any assay-labeled fitness data to use as training data for task-specific model supervision (**Figure 5**, **Supplementary Data 5**). By eliminating the reliance on any initial data collection, inverse folding has the potential to accelerate entire evolutionary campaigns.

**Figure 5:**
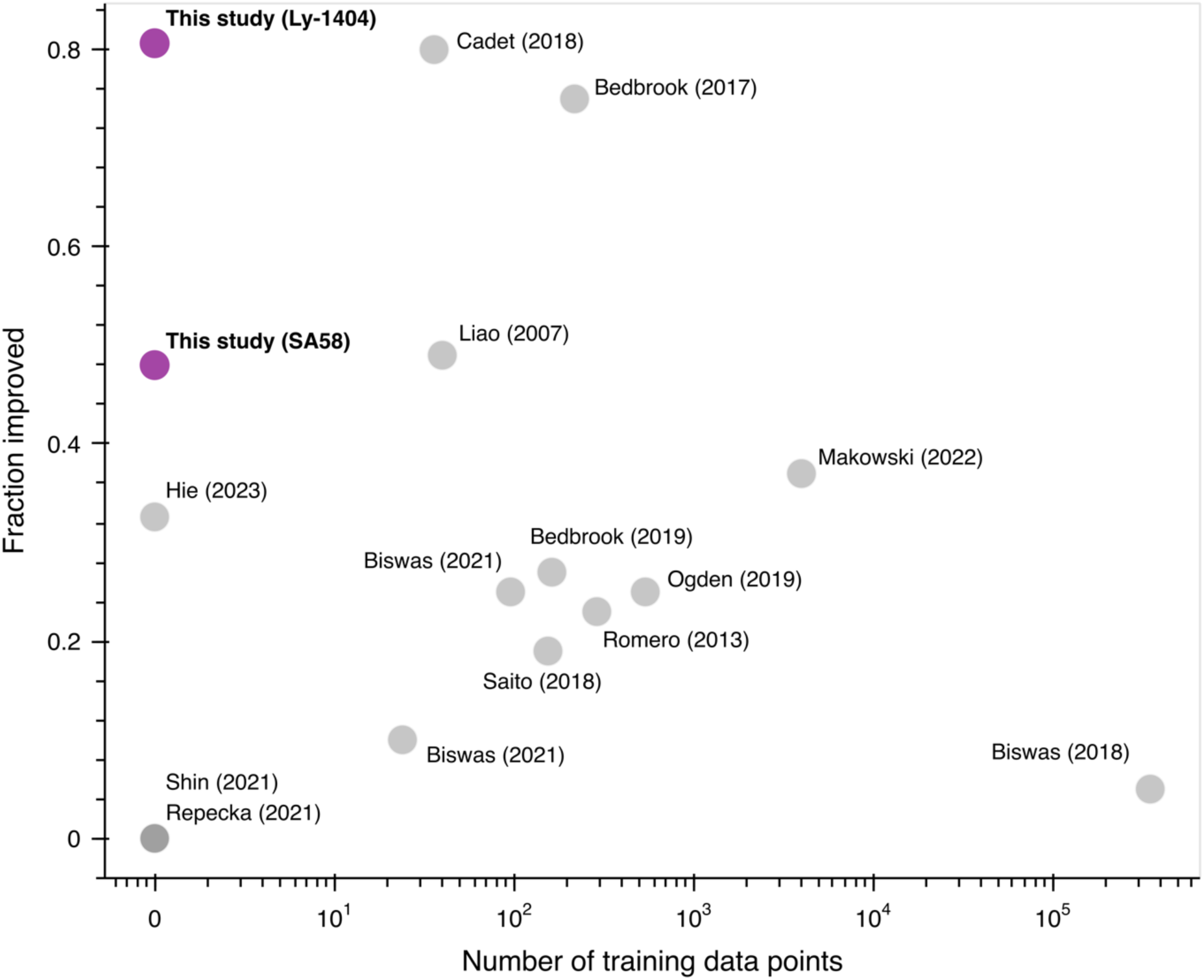
Comparison to other machine learning-guided directed evolution methods. ‘Fraction improved’ refers to the hit rate of variants tested which are improved relative to a wildtype protein used as a starting point for directed evolution or a reference protein used as a design template. Higher hit rates indicate more efficient experimental exploration. Inverse folding achieves the highest hit rate with the lowest number of assay-labeled training data points to-date^8,16–28^.

Computational methods like the one we propose have the opportunity to democratize protein engineering efforts. Not only is our approach more efficient than conventional resource-intensive techniques that experimentally test the effects of all single-residue changes on biochemical functions like binding affinity, but consequently it enables directed evolution based on properties that are not easily measured at scale or that are incompatible with high-throughput screening. Overcoming these limitations, we anticipate our structure-based paradigm will be useful for evolving proteins across many domains.

## Methods

### Inverse folding model description and scoring of sequences

As input to the inverse folding model, we provide a protein structure 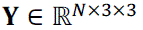, where *N* is the number of amino acids, and each amino acid is featurized by the three-dimensional physical coordinates of all three atoms in the protein backbone: the α-carbon, β-carbon, and nitrogen atoms in the protein backbone (hence the dimensionality *N* × 3 × 3). The inverse folding model learns the probability distribution *p* of a protein sequence 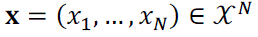 (where *X* is the alphabet of amino acids) given a structure **Y** via the chain rule of probability

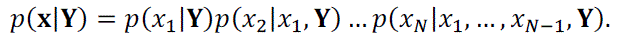

The probability distribution at each position is defined over *X*, such that it is a 20-dimensional vector with all constituent entries summing to 1.

Thus, for a given sequence 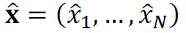 and its corresponding given structure 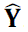, we can score the probability of 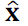 folding into **Y** under the inverse folding model by computing the value of 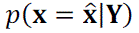, which we can do autoregressively as

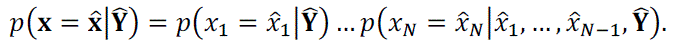

This is evaluated output is a likelihood between 0 and 1, inclusive. The computed score 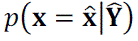 is used as prediction for “fitness” (e.g., binding affinity or enzymatic activity). Importantly, the inverse folding model does not have any explicit access to “fitness” during either training or evaluation, which we refer to as “zero shot” fitness prediction.

We use the inverse folding model checkpoint of ESM-IF1 GVP-Transformer as of April 10, 2022^11^.

### Diverse proteins benchmarking experiment with scanning mutagenesis data

We analyzed the effectiveness of using the inverse folding language model, ESM-IF1 model to identify high fitness variants from protein mutational scans as a proxy for the ability to guide evolution without explicitly modeling a protein’s function. We also compared its performance to ESM-1v, a sequence-only general protein language model. To do so, we used all deep mutational scanning (DMS) datasets from the benchmarking study by Livesey and Marsh^29^ profiling over 100 variants and reported to have 90% or higher coverage of DMS results across the corresponding curated PDB structure (**Supplementary Table 1**). From this set of 12 proteins, Cas9 was excluded because its sequence length was larger than the maximum allowable length of 1024 amino acids by ESM-1v and ccdB was excluded because the experimental values were discretized within a small range. For each of the 10 mutagenesis datasets, all the sequence likelihood of all variants with coverage in the structure were determined using inverse folding. For ESM-1v, the average masked marginals likelihood score across all five models in the ESM-1v group was used. The experimental data distribution was binarized for high-fitness classification using a percentile-based threshold. The enrichment of high fitness variants was then determined by using the metric of fraction high fitness as defined by the fraction of the top 10 model-predicted variants with experimental values above the high fitness threshold. The analysis was performed at three different percentile thresholds, top 5^th^ percentile (95^th^ percentile), top 10^th^ percentile (90^th^ percentile), and top 20^th^ percentile (80^th^ percentile), to determine sensitivity of the result based on the stringency of the selected cutoff parameter.

### Benchmarking of antibody mutagenesis

We use three antibody mutagenesis datasets^50,51^ to benchmark the performance of modeling variant effects on antibody binding using inverse folding against two sequence-only methods, ESM-1v^49^ and abYsis^55^. Variant sequences were scored using the inverse folding model with three different forms of structure input: i) variable region of mutated antibody chain only ii) variable regions of both antibody chains iii) variable regions of both antibody chains in complex with antigen. The autoregressive scoring of sequences with inverse folding enables evaluation of sequences with multiple mutations. The Spearman correlation was determined between the log likelihood scores across all sequences and corresponding reported experimental binding measurements: –log(*K*_D_) for CR9114 and CR6261; log(binding enrichment) g6.31. The following structures were used for input backbone coordinates of the VH, VL, and antigen: PDB 4FQI^52^, CR9114-H5; PDB 3GBN^53^, CR6261-H1; PDB 2FJG, g6.31-VEGF.

ESM-1v and abYsis were scored using the variant sequence of the antibody variable region. For variants with multiple mutations, the average effect of all mutant amino acids in the sequence was considered, namely

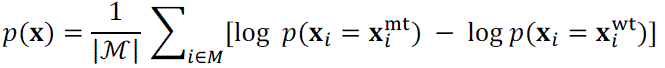

where ℳ is defined as the set of all mutations in the input sequence **X**. For ab**Y**sis, individual mutation likelihoods were determined using the frequency of amino acids at each position based on multiple sequence alignment provided by the webtool version 3.4.1 (http://www.abysis.org/abysis/index.html). We aligned VH and VL protein sequences using the default settings provided in the ‘Annotate’ tool, with the database of ‘Homo sapiens’ sequences as of April 1, 2023.

### Acquisition of antibody amino acid substitutions using inverse folding

We select amino acid substitutions recommended by the inverse folding model to test in our directed evolution campaigns for Ly-1404 and SA58. For a given wild-type antibody variable region sequence, 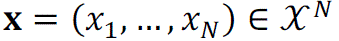, where *X* is the set of amino acids and *N* is the sequence length, we score all possible single amino acid substitutions against a corresponding structure of the variable regions of both antibody chains in complex with the RBD of SARS-CoV-2 Spike protein, 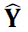 by computing 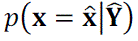. Protein structures used are reported in Supplementary Table 1. We then select the set of top ten predicted single amino acid substitutions at unique residues in each antibody variable region to test in the first round of evolution.

After testing individual amino acid mutations in a pseudovirus neutralization screen, in Round 2, beneficial mutations (defined as IC_50_ fold-change > 1.1) were combined to assess the combinatorial effects and potential for further neutralization improvement. We tested up to four combinations of single amino acid mutations on each chain (two total mutations to the antibody). We also used the inverse folding model to score a library of all possible combinations of the beneficial mutations to an antibody chain (For example, VH Ly-1404 has 8 beneficial mutations resulting in 255 total candidate sequences), and selected the top five scoring designs (or less if there were a fewer number of total possible combinations). Lastly, we tested a maximum of two variants consisting of the best single-chain designs together. In total, 31 variants were tested for Ly-1404 and 25 variants were tested for SA58.

### Antibody cloning

We cloned the antibody sequences into the CMV/R plasmid backbone for expression under a CMV promoter. The heavy chain or light chain sequence was cloned between the CMV promoter and the bGH poly(A) signal sequence of the CMV/R plasmid to facilitate improved protein expression. Variable regions were cloned into the human IgG1 backbone; Ly-1404 variants were cloned with a lambda light chain, whereas variants of SA58 were cloned with a kappa light chain. The vector for both heavy and light chain sequences also contained the HVM06_Mouse (UniProt: <u>P01750</u>) Ig heavy chain V region 102 signal peptide (MGWSCIILFLVATATGVHS) to allow for protein secretion and purification from the supernatant. VH and VL segments were ordered as gene blocks from Integrated DNA Technologies and were cloned into linearized CMV/R backbones with 5× In-Fusion HD Enzyme Premix (Takara Bio).

### Antigen cloning

RBD sequences were cloned into a pADD2 vector between the rBeta-globin intron and β-globin poly(A). All RBD constructs contain an AviTag and 6×His tag. RBD sequences were based off wild-type Wuhan-Hu-1 (GenBank: <u>BCN86353.1</u>), Omicron BA.1 (GenBank: <u>UFO69279.1</u>), BQ.1.1 (GenBank: <u>OP412163.1</u>), XBB.1.5 (GenBank: <u>OP790748.1</u>).

### DNA preparation

Plasmids were transformed into Stellar competent cells (Takara Bio), and transformed cells were plated and grown at 37 °C overnight. Colonies were mini-prepped per the manufacturer’s recommendations (GeneJET, K0502, Thermo Fisher Scientific) and sequence confirmed (Sequetech) and then maxi-prepped per the manufacturer’s protocols (ZymoPure II Plasmid Maxiprep Kit, Zymo Research). Plasmids were sterile filtered using a 0.22-μm syringe filter and stored at 4 °C.

### Protein expression

All proteins were expressed in Expi293F cells (Thermo Fisher Scientific, A14527). Proteins containing a biotinylation tag (AviTag) were also expressed in the presence of a BirA enzyme, resulting in spontaneous biotinylation during protein expression. Expi293F cells were cultured in media containing 66% FreeStyle/33% Expi media (Thermo Fisher Scientific) and grown in TriForest polycarbonate shaking flasks at 37 °C in 8% carbon dioxide. The day before transfection, cells were pelleted by centrifugation and resuspended to a density of 3 × 10^6^ cells per milliliter in fresh media. The next day, cells were diluted and transfected at a density of approximately 3–4 × 10^6^ cells per milliliter. Transfection mixtures were made by adding the following components: maxi-prepped DNA, culture media and FectoPRO (Polyplus) would be added to cells to a ratio of 0.5 μg: 100 μl: 1.3 μl: 900 μl. For example, for a 100-ml transfection, 50 μg of DNA would be added to 10 ml of culture media, followed by the addition of 130 μl of FectoPRO. For antibodies, we divided the transfection DNA equally among heavy and light chains; in the previous example, 25 μg of heavy chain DNA and 25 μg of light chain DNA would be added to 10 ml of culture media. After mixing and a 10-min incubation, the example transfection cocktail would be added to 90 ml of cells. The cells were harvested 3–5 days after transfection by spinning the cultures at 10,000*g* for 10 min. Supernatants were filtered using a 0.45-μm filter.

### Antibody purification

We purified antibodies using a 5-ml MabSelect Sure PRISM column on the ÄKTA pure fast protein liquid chromatography (FPLC) instrument (Cytiva). The ÄKTA system was equilibrated with line A1 in 20 mM 4-(2-hydroxyethyl)-1-piperazineethanesulfonic acid (HEPES) pH 7.4, 150 mM sodium chloride (NaCl), line A2 in 100 mM glycine pH 2.8, line B1 in 0.5 M sodium hydroxide, Buffer line in 20 mM 4-(2-hydroxyethyl)-1-piperazineethanesulfonic acid (HEPES) pH 7.4, 150 mM sodium chloride (NaCl) and Sample lines in water. The protocol washes the column with A1, followed by loading of the sample in the Sample line until air is detected in the air sensor of the sample pumps, followed by five column volume washes with A1, elution of the sample by flowing of 20 ml of A2 directly into a 50-ml conical containing 2 ml of 1 M tris(hydroxymethyl)aminomethane (Tris) pH 8.0, followed by five column volumes of A1, B1 and A1 and then a wash step of the fraction collector with A1. We concentrated the eluted samples using 50-kDa cutoff centrifugal concentrators, followed by buffer exchange using a PD-10 column (Sephadex) that had been pre-equilibrated into 20 mM 4-(2-hydroxyethyl)-1-piperazineethanesulfonic acid (HEPES) pH 7.4, 150 mM sodium chloride (NaCl). Purified antibodies were used directly in experiments or flash-frozen and stored at −20 °C.

### Antigen purification

All RBD antigens were His-tagged and purified using HisPur Ni-NTA resin (Thermo Fisher Scientific, 88222). Cell supernatants were diluted with 1/3 volume of wash buffer (20 mM imidazole, 20 mM 4-(2-hydroxyethyl)-1-piperazineethanesulfonic acid (HEPES) pH 7.4, 150 mM sodium chloride (NaCl), and the Ni-NTA resin was added to diluted cell supernatants. For all antigens, the samples were then incubated at 4 °C while stirring overnight.

Resin/supernatant mixtures were added to chromatography columns for gravity flow purification. The resin in the column was washed with wash buffer (20 mM imidazole, 20 mM HEPES pH 7.4, 150 mM NaCl), and the proteins were eluted with 250 mM imidazole, 20 mM HEPES pH 7.4, 150 mM NaCl. Column elutions were concentrated using centrifugal concentrators at 10-kDa cutoff, followed by size-exclusion chromatography on an ÄKTA pure system (Cytiva).

ÄKTA pure FPLC with a Superdex 200 Increase (S200) gel filtration column was used for purification. Then, 1 ml of sample was injected using a 2-ml loop and run over the S200, which had been pre-equilibrated in degassed 20 mM HEPES, 150 mM NaCl before use and flash-frozen before storage at −20 °C.

### BLI binding experiments

All reactions were run on an Octet RED96 at 30 °C, and samples were run in 1× PBS with 0.1% BSA and 0.05% Tween 20 (Octet buffer). IgGs were assessed for binding to biotinylated antigens using streptavidin biosensors (Sartorius/ForteBio). Antigen was loaded at a concentration of 200nM. Tips were then washed and baselined in wells containing only Octet buffer. Samples were then associated in wells containing IgG at 100 nM concentration. A control well with loaded antigen but that was associated in a well containing only 200 μl of Octet buffer was used as a baseline subtraction for data analysis. Association and dissociation binding curves were fit in Octet System Data Analysis Software version 9.0.0.15 using a 1:2 bivalent model for IgGs to determine apparent *K*_d_. Fold-change in apparent *K*_d_ were determined by computing the ratio of wildtype *K*_d_ to variant *K*_d_. Averages of *K*_d_ fold-change values from at least two independent experiments are reported to two significant figures in **Supplementary Data 2**. To estimate measurement error, we computed the standard deviation for each antibody−antigen *K*_d_ pair.

### Polyspecificity Particle assay

Polyspecificity reagent (PSR) was obtained as described by Xu et al^63^. Soluble membrane proteins were isolated from homogenized and sonicated Expi 293F cells followed by biotinylation with Sulfo-NHC-SS-Biotin (Thermo Fisher Scientific, 21331) and stored in PBS at −80 °C. The PolySpecificity Particle (PSP) assay was performed as described in Makowski et al.^64^. Protein A magnetic beads (Invitrogen, 10001D) were washed three times in PBSB (PBS with 1 mg ml^−1^ BSA) and diluted to 54 μg ml^−1^ in PBSB. Then, 30 μl of the solution containing the beads was incubated with 85 μl of antibodies at 15 µg ml^−1^ overnight at 4 °C with rocking. The coated beads were then washed twice with PBSB using a magnetic plate stand (Invitrogen, 12027) and resuspended in PBSB. We then incubated 50 μl of 0.1 mg ml^−1^ PSR with the washed beads at 4 °C with rocking for 20 min. Beads were then washed with PBSB and incubated with 0.001× streptavidin-APC (BioLegend, 405207) and 0.001× goat anti-human Fab fragment FITC (Jackson ImmunoResearch, 109-097-003) at 4 °C with rocking for 15 min. Beads were then washed and resuspended with PBSB. Beads were profiled via flow cytometry using a Sony SH800 cell sorter. Data analysis was performed with FlowJo software version 10.9.0 to obtain median fluorescence intensity (MFI) values, which are reported for each antibody across three or more replicate wells. Elotuzumab (Fisher Scientific) and ixekizumab (Fisher Scientific) are also included in each assay as controls.

### Lentivirus production

We produced SARS-CoV-2 Spike (Wuhan-Hu-1, BA.1, and BQ.1.1 variants) pseudotyped lentiviral particles. Viral transfections were done in HEK293T cells (American Type Culture Collection, CRL-3216) using BioT (BioLand) transfection reagent. Six million cells were seeded in D10 media (DMEM + additives: 10% FBS, L-glutamate, penicillin, streptomycin and 10 mM HEPES) in 10-cm plates one day before transfection. A five-plasmid system was used for viral production, as described in Crawford et al^65^. The Spike vector contained the 21-amino-acid truncated form of the SARS-CoV-2 Spike sequence from the Wuhan-Hu-1 strain of SARS-CoV-2 (GenBank: <u>BCN86353.1</u>), BA.1 variant of concern (GenBank: <u>OL672836.1</u>), or BQ.1.1 variant of concern (GenBank: <u>OP412163.1</u>. The other viral plasmids, used as previously described^65^, are pHAGE-Luc2-IRS-ZsGreen (NR-52516), HDM-Hgpm2 (NR-52517), pRC-CMV-Rev1b (NR-52519) and HDM-tat1b (NR-52518). These plasmids were added to D10 medium in the following ratios: 10 μg pHAGE-Luc2-IRS-ZsGreen, 3.4 μg FL Spike, 2.2 μg HDM-Hgpm2, 2.2 μg HDM-Tat1b and 2.2 μg pRC-CMV-Rev1b in a final volume of 1,000 μl.

After adding plasmids to medium, we added 30 μl of BioT to form transfection complexes. Transfection reactions were incubated for 10 min at room temperature, and then 9 ml of medium was added slowly. The resultant 10 ml was added to plated HEK cells from which the medium had been removed. Culture medium was removed 24 h after transfection and replaced with fresh D10 medium. Viral supernatants were harvested 72 h after transfection by spinning at 300*g* for 5 min, followed by filtering through a 0.45-μm filter. Viral stocks were aliquoted and stored at −80 °C.

### Pseudovirus neutralization

The target cells used for infection in SARS-CoV-2 pseudovirus neutralization assays are from a HeLa cell line stably overexpressing human angiotensin-converting enzyme 2 (ACE2) as well as the protease known to process SARS-CoV-2: transmembrane serine protease 2 (TMPRSS2). Production of this cell line is described in detail by Rogers et al^66^. with the addition of stable TMPRSS2 incorporation. ACE2/TMPRSS2/HeLa cells were plated 1 day before infection at 8,000 cells per well. Ninety-six-well, white-walled, white-bottom plates were used for neutralization assays (Thermo Fisher Scientific).

On the day of the assay, purified IgGs in 1× PBS were made into D10 medium (DMEM + additives: 10% FBS, L-glutamate, penicillin, streptomycin and 10 mM HEPES). A virus mixture was made containing the virus of interest (for example, SARS-CoV-2) and D10 media. Virus dilutions into media were selected such that a suitable signal would be obtained in the virus-only wells. A suitable signal was selected such that the virus-only wells would achieve a luminescence of at least >1,000,000 relative light units (RLU). Then, 60 μl of this virus mixture was added to each of the antibody dilutions to make a final volume of 120 μl in each well. Virus-only wells were made, which contained 60 μl of D10 and 60 μl of virus mixture. Cells-only wells were made, which contained 120 μl of D10 media.

The antibody/virus mixture was left to incubate for 1 h at 37 °C. After incubation, the medium was removed from the cells on the plates made one day prior. This was replaced with 100 μl of antibody/virus dilutions and incubated at 37 °C for approximately 48 h. Infectivity readout was performed by measuring luciferase levels. Medium was removed from all wells, and cells were lysed by the addition of 100 μl of BriteLite assay readout solution (PerkinElmer) into each well. Luminescence values were measured using an Infinite 200 PRO Microplate Reader (Tecan) using i-control version 2.0 software (Tecan) after shaking for 30 sec. Each plate was normalized by averaging the cells-only (0% infection) and virus-only (100% infection) wells. Neutralization titer was defined as the sample dilution at which the RLU was decreased by 50% as compared with the RLU of virus-only control wells after subtraction of background RLUs in wells containing cells only. Normalized values were fitted with a three-parameter nonlinear regression inhibitor curve in GraphPad Prism 9.1.0 to determine the half-maximal inhibitory concentration (IC_50_) and are reported in **Supplementary Data 1**. Neutralization assays were performed in biological duplicates with technical duplicates.

### Computing frequency of changes to antibody protein sequences

We computed the frequency of residues involved in affinity-enhancing substitutions using the abYsis webtool, which also computes the frequency of amino acids at each position based on a multiple sequence alignment. We aligned VH and VL protein sequences using the default settings provided in the ‘Annotate’ tool, using the database of ‘All’ sequences as of April 1, 2023. We also used the Kabat region definition provided by abYsis webtool version 3.4.1 to annotate the framework regions and CDRs within the VH and VL sequences which are reported in **Supplementary Table 2**.

### Comparing efficiency of machine learning-guided directed evolution methods

To compare inverse folding against other machine learning methods for protein evolution, we compared the fraction of variants tested in the protein engineering campaign to the number of assay-labeled training data points used to inform the predictions. Data was sourced from Biswas et al.^17^ and made contemporaneous by the addition of recently published studies as indicated in **Supplementary Data 5**. The fraction improved, or hit rate, refers to experimentally tested predictions which have improved functional activity relative to either a wildtype protein that is used as a starting point for directed evolution or the protein used as a reference template for design.

## Acknowledgments

We would like to thank D. Xu, S. Kim, and the members of the Kim lab for helpful discussions on this project. We are also grateful for assistance from D. Xu with protein graphics. V.R.S acknowledges the support of the Stanford University Medical Scientist Training Program grants (T32-GM007365 and T32-GM145402). V.R.S. and T.U.J.B. are both supported by the Sarafan ChEM-H Chemistry/Biology Interface Training Program. T.U.J.B. is also supported by the Knight-Hennessy Graduate Scholarship Fund and a CIHR Doctoral Foreign Study Award (FRN:170770). B.L.H is supported by the Stanford Science Fellows Program. This work was supported by the Virginia & D.K. Ludwig Fund for Cancer Research and the Chan Zuckerberg Biohub.

## Author contributions

Conceptualization, methodology, interpretation: V.R.S., B.L.H., P.S.K.; Computational experiments and software development: V.R.S.; Antibody and antigen cloning, expression, and purification: V.R.S., T.U.J.B.; Lentivirus production and pseudovirus neutralization: T.U.J.B; Binding assays: V.R.S.; Writing (original draft): V.R.S with assistance from B.L.H and P.S.K.; Writing (final draft): all authors

## Supplementary Figures, Tables, Information, & Data

**Supplementary Table 1:** List of proteins, protein structures, and assay information for deep mutational scanning experiments

**Supplementary Table 2**: Analysis of neutralization-enhancing mutations

**Supplementary Information**: Antibody sequences

**Supplementary Data 1**: Neutralization data with IC_50_ values of evolved antibodies across both evolutionary campaigns

**Supplementary Data 2:** Binding data with IgG *K*_D_ values of evolved antibodies

**Supplementary Data 3**: Antibody variant prediction benchmarking results

**Supplementary Data 4**: MFI values for polyspecificity experiments

**Supplementary Data 5**: Efficiency comparison of machine learning-guided directed evolution methods

**Supplementary Figure 1:**
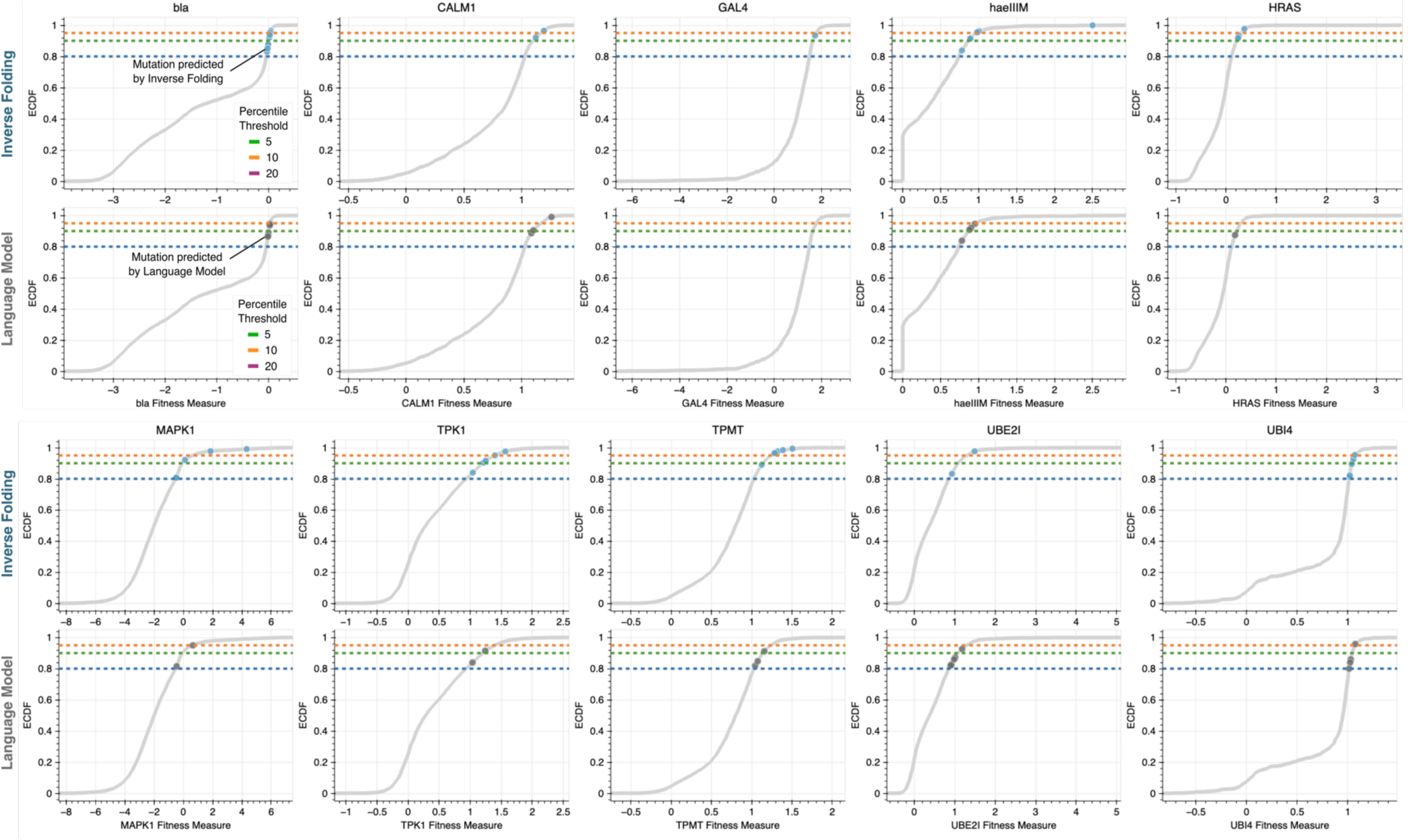
Inverse folding identifies high fitness variants across proteins with diverse functions. In addition to higher hit rates of high fitness variants, inverse folding generally identifies variants with greater magnitude of improvements in fitness. The top ten predicted variants with experimental fitness values ranking in the 20^th^ percentile of all variants profiled in the deep mutational screen are shown. The grey curve shows the empirical cumulative distribution function (ECDF) of all experimental fitness values determined in the screen. The dotted lines correspond to the three percentile-based thresholds used in the sensitivity analysis (**Figure 1d)** to classify high fitness variants. bla, Beta-lactamase TEM; CALM1, Calmodulin-1; haeIIIM, Type II methyltransferase M.HaeIII; HRAS, GTPase HRas; MAPK1, Mitogen-activated protein kinase; TMPT, Thiopurine S-methyltransferase; TPK1, Thiamin pyrophosphokinase 1; UBI4, Polyubiquitin; UBE2I, SUMO-conjugating enzyme UBC9

**Supplementary Figure 2:**
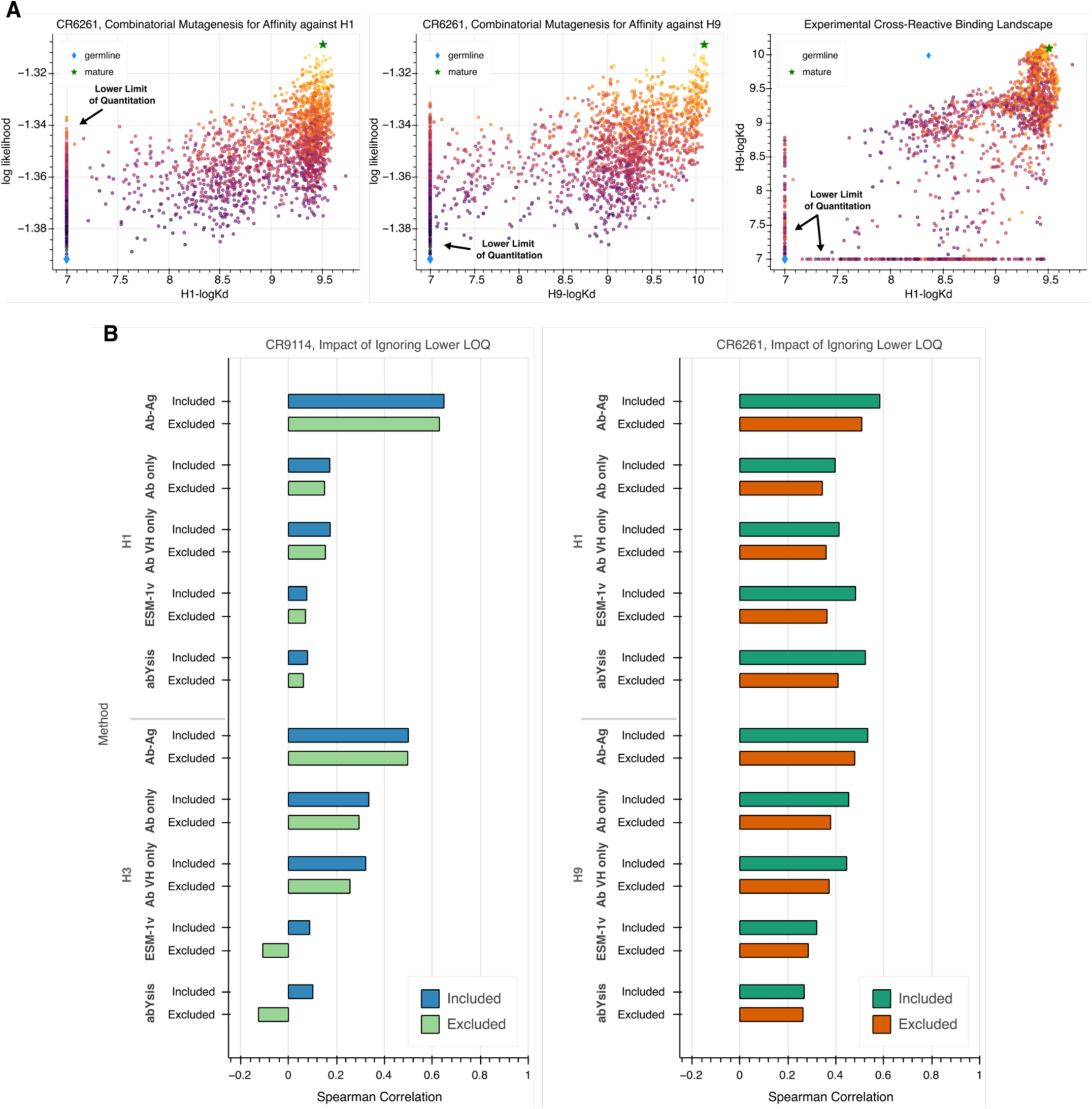
Impact of lower limit of quantitation of binding assay on predictive performance (A) Scatter plots showing CR6261 variant sequences scored with inverse folding compared to experimental binding data and inclusive of the assay’s lower limit of quantitation, which is omitted for visualization in **Figure 3b**. **(B)** Comparative bar plots showing the impact of removing sequences with experimental measurements bounded artificially by the assay to dataset-wide correlation. While Spearman correlations shown in Figure 3a are computed without any modification to the data, trends in prediction and comparison among modeling methods are robust to filtering sequences affected by this assay artifact.

**Supplementary Figure 3:**
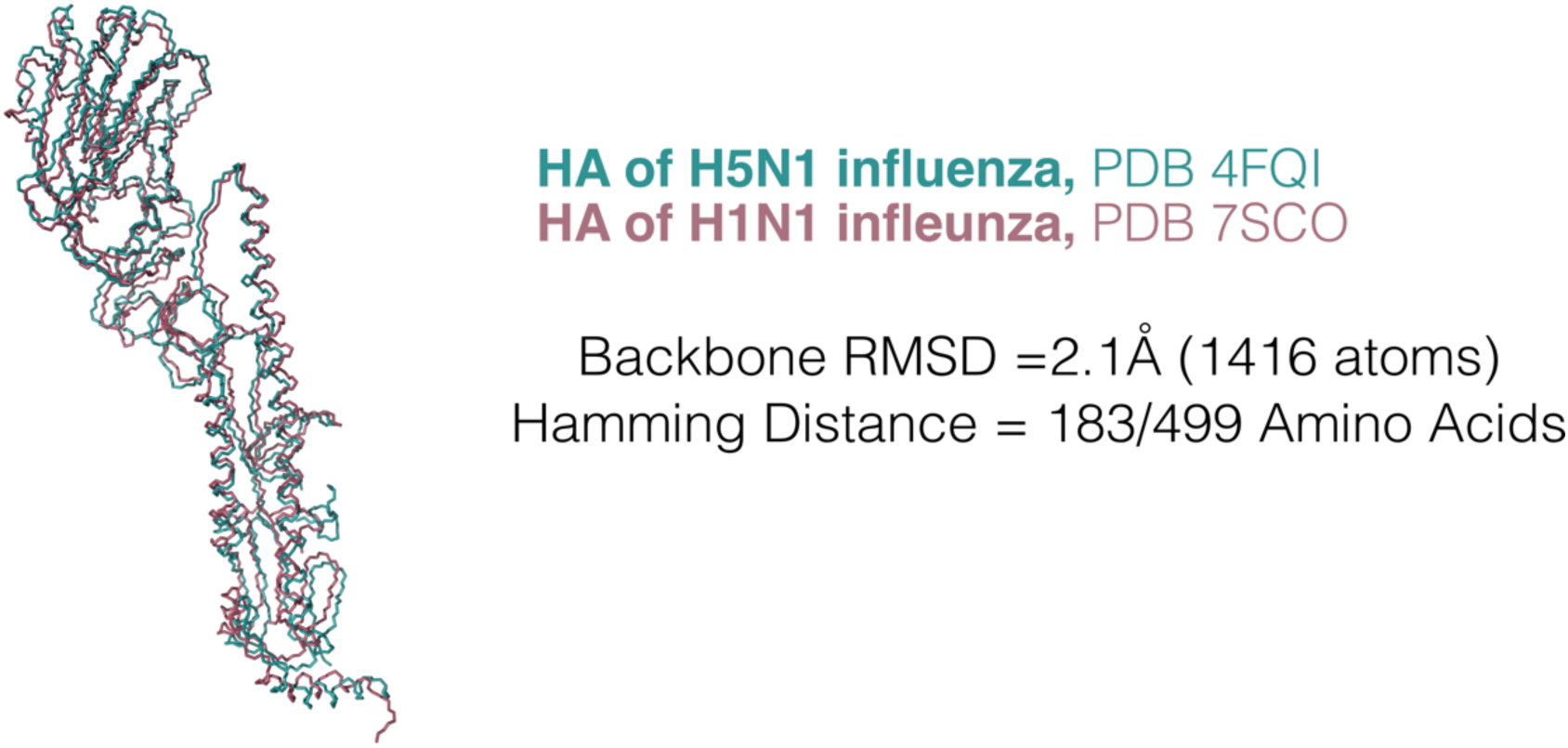
Structural and sequence similarity of H5 and H1. For cross-reactive antibodies, inclusion of the antigen structure is informative even for predicting binding to a different antigen. In Figure 3a, we report a correlation of 0.65 between inverse folding log likelihoods of CR9114 variants and experimental affinity measurements to H1 despite using a structure solved with CR9114 in complex with H5. Inverse folding uses both the protein sequence and backbone structure coordinates as input. Across both HA subunits, H5 and H1 have considerable sequence differences and a 2.1 Å root mean square deviation (RMSD) across the entire protein backbone.

**Supplementary Figure 4:**
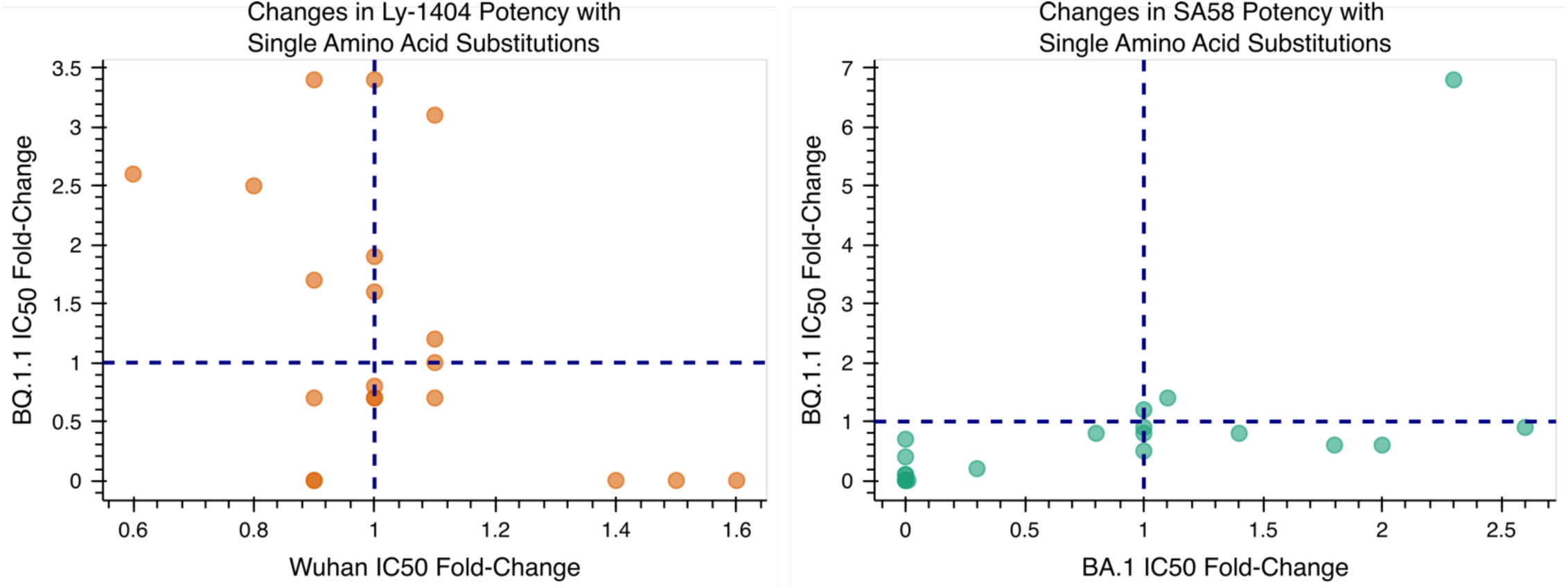
Functional diversity of inverse folding-recommended mutations. Among the 20 single amino acid substitutions tested for Ly-1404, 14 of 20 = 70% improve neutralization against at least one of the two strains tested. Similarly, 7 of 20 = 35% of the single amino acid substitutions tested for SA58 improve neutralization. While some variants improve function against both pseudovirus strains, others overwhelmingly only improve against one. This suggests that focusing sequence exploration to structurally compatible mutations does not compromise functional diversity.

**Supplementary Figure 5:**
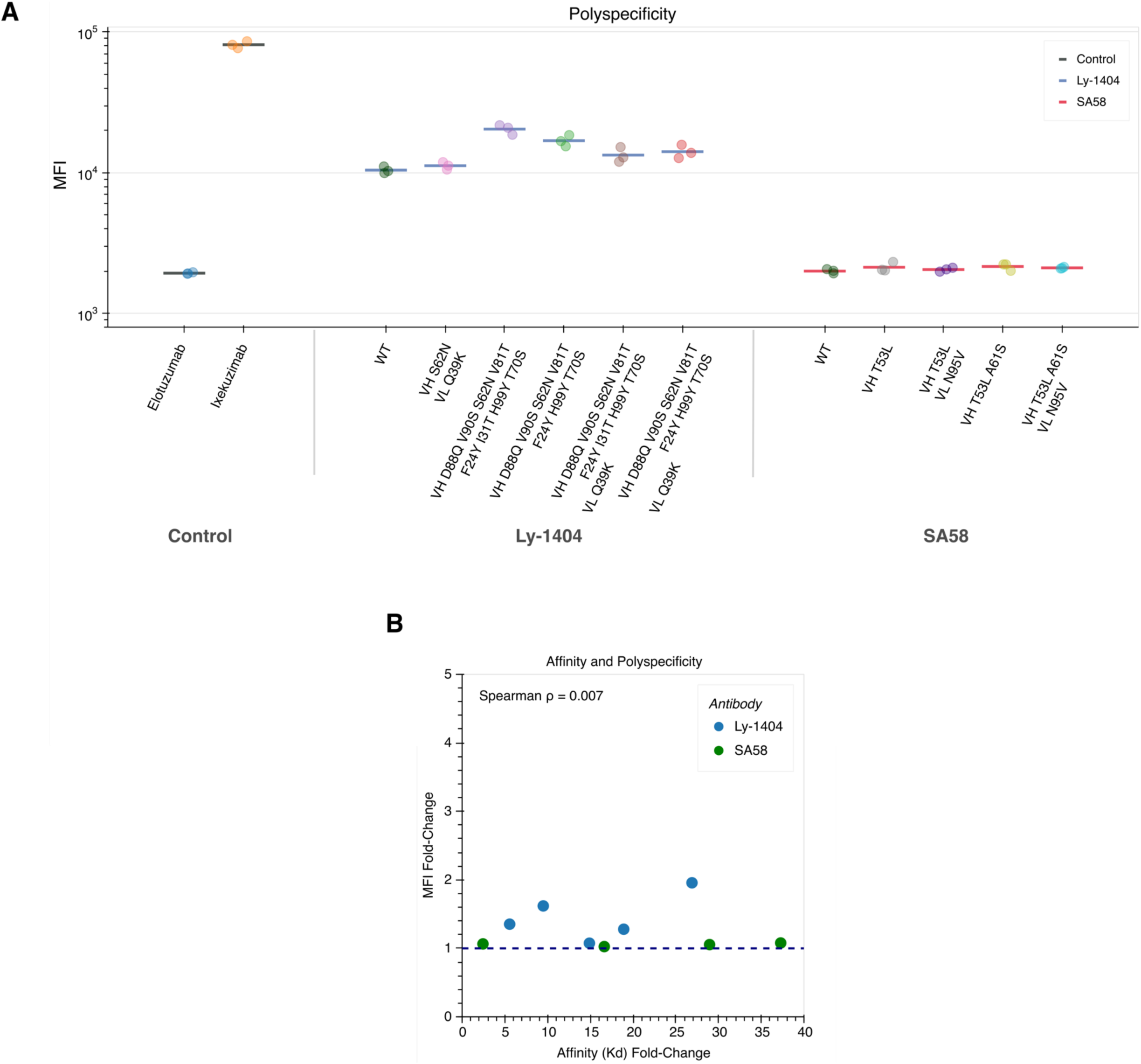
Polyspecificity of evolved antibodies (A) The median fluorescence intensity (MFI) signal obtained from flow cytometry is shown for several evolved antibodies with improved affinity and compared to two clinical monoclonal antibodies with high and low polyspecificity used to define a clinically viable range. **(B)** Fold-change in polyspecificity signal is plotted against fold-change in affinity as IgG against BQ.1.1 for Ly-1404 and XBB.1.5 for SA58. There is no correlation between the improvements in on-target improvements in affinity and off-target nonspecific changes in polyspecificity (Spearman π = 0.007).

**Supplementary Figure 6:**
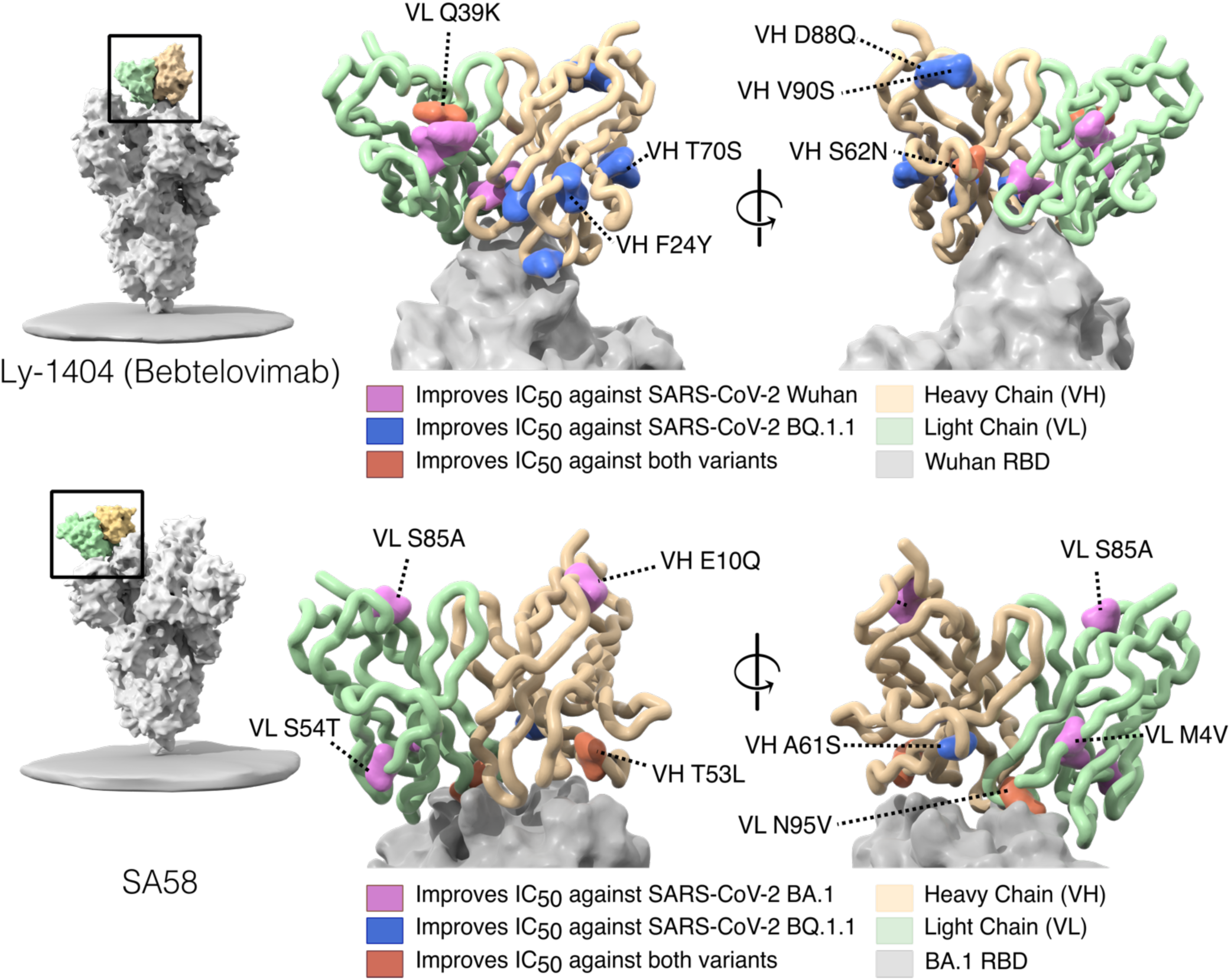
Mapping neutralization-enhancing substitutions. Neutralization-enhancing mutations are labeled on the structure of the wild-type antibody in complex with the RBD of SARS-CoV-2 spike protein (Ly-1404: PDB 7mmo; SA58: PDB 7y0w). Notably, several mutations are identified to have significant beneficial impacts on binding neutralization and affinity (**Supplementary Data 1 & 2)** despite located away from the binding interface.

**Supplementary Table 1.** Summary of the DMS datasets used in this analysis, including functional assay, method of mutagenesis, and structure used for inverse folding scoring. We also note the specific DMS assay from each study we use for calculating correlation with inverse folding log likelihoods.

| Protein(s)<br>(Uniprot ID) | Organism | Functional Assay | Mutagenesis Method | Utilized assay | PDB Structure | Total coverage of DMS (%) | Access date* | Reference |
| --- | --- | --- | --- | --- | --- | --- | --- | --- |
| UBE2I (P63279) | Human | POPCode, a variant of multiple-site directed mutagenesis. | Competitive growth assay in yeast. | score | 5F6E chain A | 100 | 12/10/2018 | (Weile *et al*, 2017) |
| TPK1 (Q9H3S3) |  |  |  | score | 3S4Y chain A | 92.46 |  |  |
| CALM1 (P0DP23) |  |  |  | score | 5V03 chain R | 100 |  |  |
| HRas (P01112) | Human | Systematic site-directed mutagenesis. | Two-hybrid assay. | unregulated | 2CE2 chain X | 100 | 12/10/2018 | (Bandaru *et al*, 2017) |
| MAPK1 (P28482) | Human | Systematic site-directed mutagenesis. | Competitive growth assay. | VRT | 4ZZN chain A | 99.44 | 12/10/2018 | (Brenan *et al*, 2016) |
| TPMT (P51580) | Human | Systematic site-directed mutagenesis. | Fluorescence of a GFP fusion protein. | score | 2BZG chain A | 92.9 | 12/10/2018 | (Matreyek *et al*, 2018) |
| UBI4(b) (P0CG63) | Yeast | Site directed mutagenesis by cassette ligation. | Fluorescence activated cell sorting (FACS). | Relative_E1-activity_limiting | 4Q5E chain B | 100 | 12/10/2018 | (Roscoe & Bolon, 2014) |
| GAL4 (P04386) | Yeast | Systematic site-directed mutagenesis. | Two-hybrid assay. | Nonselection_24 | 3COQ chain B | 90.64 | 12/10/2018 | (Kitzman *et al*, 2015) |
| bla(b) (P62593) | E. coli | Systematic site-directed mutagenesis. | Antibiotic resistance. | Ampicillin_2500 | 1M40 chain A | 100 | 12/10/2018 | (Stiffler *et al*, 2015) |
| haeIIIM (P20589) | H. aegyptius | Random mutagenesis. | Competitive growth assay. | DMS_G3 | 3UBT chain B | 99.37 | 12/10/2018 | (Rockah-Shmuel *et al*, 2015) |

\*Access date is as reported in *Livesey & Marsh*, 2020 study from which these data were sourced and this table was adapted

**Supplementary Table 2.** Single amino acid substitutions with beneficial effects on neutralization are reported alongside the region of the variable domain they are located within, as well as the wild-type and mutant amino acid frequencies in observed human antibody sequences.

**Ly-1404**
| Chain Mutated | Design | Region | WT Amino Acid Frequency | Mutant Amino Acid Frequency |
| --- | --- | --- | --- | --- |
| HC | D88Q | HFR3 | 0.03333 | 0.00382 |
| HC | V90S | HFR3 | 0.03316 | 0.05155 |
| HC | S62N | CDR-H2 | 0.13159 | 0.16299 |
| HC | V81T | HFR3 | 0.03432 | 0.00205 |
| HC | F24Y | HFR1 | 0.01738 | 0.00002 |
| HC | I31T | CDR-H1 | 0.00933 | 0.09048 |
| HC | H99Y | HFR3 | 0.01593 | 0.00138 |
| HC | T70S | HFR3 | 0.88405 | 0.06153 |
| HC | I105L | CDR-H3 | 0.02764 | 0.05760 |
| LC | A98I | CDR-L3 | 0.02297 | 0.03198 |
| LC | Q39K | LFR2 | 0.92316 | 0.00238 |
| LC | T5Q | LFR1 | 0.89340 | 0.00933 |
| LC | K47E | LFR2 | 0.52285 | 0.01490 |
| LC | M49L | LFR2 | 0.05585 | 0.77076 |

**SA58**
| Chain Mutated | Design | Region | WT Amino Acid Frequency | Mutant Amino Acid Frequency |
| --- | --- | --- | --- | --- |
| HC | T53L | CDR-H2 | 0.03814 | 0.00963 |
| HC | A61S | CDR-H2 | 0.59797 | 0.13159 |
| HC | E10Q | HFR1 | 0.24182 | 0.01366 |
| LC | N95V | CDR-L3 | 0.13399 | 0.00685 |
| LC | S85A | LFR3 | 0.01109 | 0.00698 |
| LC | S54T | CDR-L2 | 0.65138 | 0.05372 |
| LC | M4V | LFR1 | 0.29424 | 0.03348 |

